# DENSS-Multiple: A Structure Reconstruction Method using Multiple Contrast Variation of Small-Angle Neutron Scattering Based on the DENSS Algorithm

**DOI:** 10.1101/2022.03.01.478978

**Authors:** Jacob Sumner, Shuo Qian

## Abstract

Small-angle neutron scattering (SANS) provides easily manipulable neutron contrast variation that has been widely exploited to study the structure and function of biological macromolecules and their complex in solution. Here, we developed a method called DENSS-Multiple based on the DENSS (DENsity from Solution Scattering) algorithm to provide ab initio structure reconstruction with SANS contrast variation data. This new tool can exploit additional information in different SANS contrasts for improved ab initio reconstruction results.

## Introduction

Small-angle scattering (SAS) with X-rays or neutrons has been widely used for understanding the structure of a wide range of materials. SAS’s ability to observe samples, especially biomolecules in their solution state, is valuable for providing insight on the structure-function relationship in a biologically relevant condition.

In contrast to X-ray photons, which scatter off an atom’s electron cloud to provide a structural depiction of electron density, neutrons scatter off nuclei, and the scattering power is a property of nuclear interactions. Owing to compositional differences between different classes of biomolecules such as proteins, nucleic acids, and lipids, each biomolecular class has a distinct scattering length density (SLD) in neutron scattering compared with X-ray scattering^1^. The notable difference between hydrogen isotopes, protium and deuterium, allows manipulation of the small-angle neutron scattering (SANS) contrast for the molecule via isotope replacement on hydrogen-rich biomolecules. In SANS studies, a neutron contrast variation (NCV) series takes advantage of the adjustable SLD difference between components in a sample by tuning the solution buffer D_2_O:H_2_O ratio to better elucidate the biomolecular structure.^1,2^ For example, contrast matching can be used to match the SLD of specific components or phases in a complex system to provide the structure of a selected component.^1–3^ Previously, the MONSA software of ATSAS was widely used in the SANS community for multiphase structures with a bead-modeling approach.^4^ DENSS-Multiple allows for multiphase structures to be resolved without the a priori inclusion of the number of phases and phase volume, which introduce an inherent bias to the output.

### DENSS-Multiple

Recently, the DENsity from Solution Scattering (DENSS) ab initio iterative structure factor retrieval algorithm was developed for small-angle X-ray scattering (SAXS) (and implicitly SANS) data. We developed the DENSS-Multiple method, which is based on the DENSS method, for reconstructing a single synthesized structure from multiple SANS data sets in a contrast-variation series.^5^ DENSS-Multiple requires minimal a priori information of the biomolecular system. With parallel iterative reconstructions from multiple data sets, DENSS-Multiple can improve the overall structure and internal SLD compared with results from a single scattering profile, thereby taking advantage of the distinct shape and density information contained in each SANS data set.

The required inputs for DENSS-Multiple are NCV data sets, which usually consist of more than one scattering curve, and the solution background SLD for each contrast, which is approximated from the H_2_O:D_2_O ratios. The process begins with a common starting model density generated randomly (Figure 1) in a 3D array. The modeled 3D density is duplicated with the number of contrasts. For each contrast, the copied 3D density is then corrected for the background SLD of each contrast (i.e., the SLD of the particular H_2_O:D_2_O buffer concentration as a priori information) by subtracting the SLD uniformly from the starting model density. This step creates different corrected contrasts, Δρ = ρ_sample_ -ρ_buffer_, in vacuo for generating the structure factors in the reciprocal space. Data from all contrast go through the steps of the phase retrieval algorithm as parallel processes (Figure 1), including fast Fourier transform (FFT) and structure-factor scaling. Once the structure factors are rescaled according to each scattering profile, they are transformed into real-space model densities using inverse FFT. The real-space constraints from the DENSS algorithm (e.g., solvent flattening) can be independently applied to each contrast by the user. The resultant model density from each contrast is then rectified with the addition of the background SLDs within the support boundary, which means the reconstruction result effectively remains in vacuo and can be combined into a single overall model density by averaging all contrasts for the next iteration.

**Figure 1:**
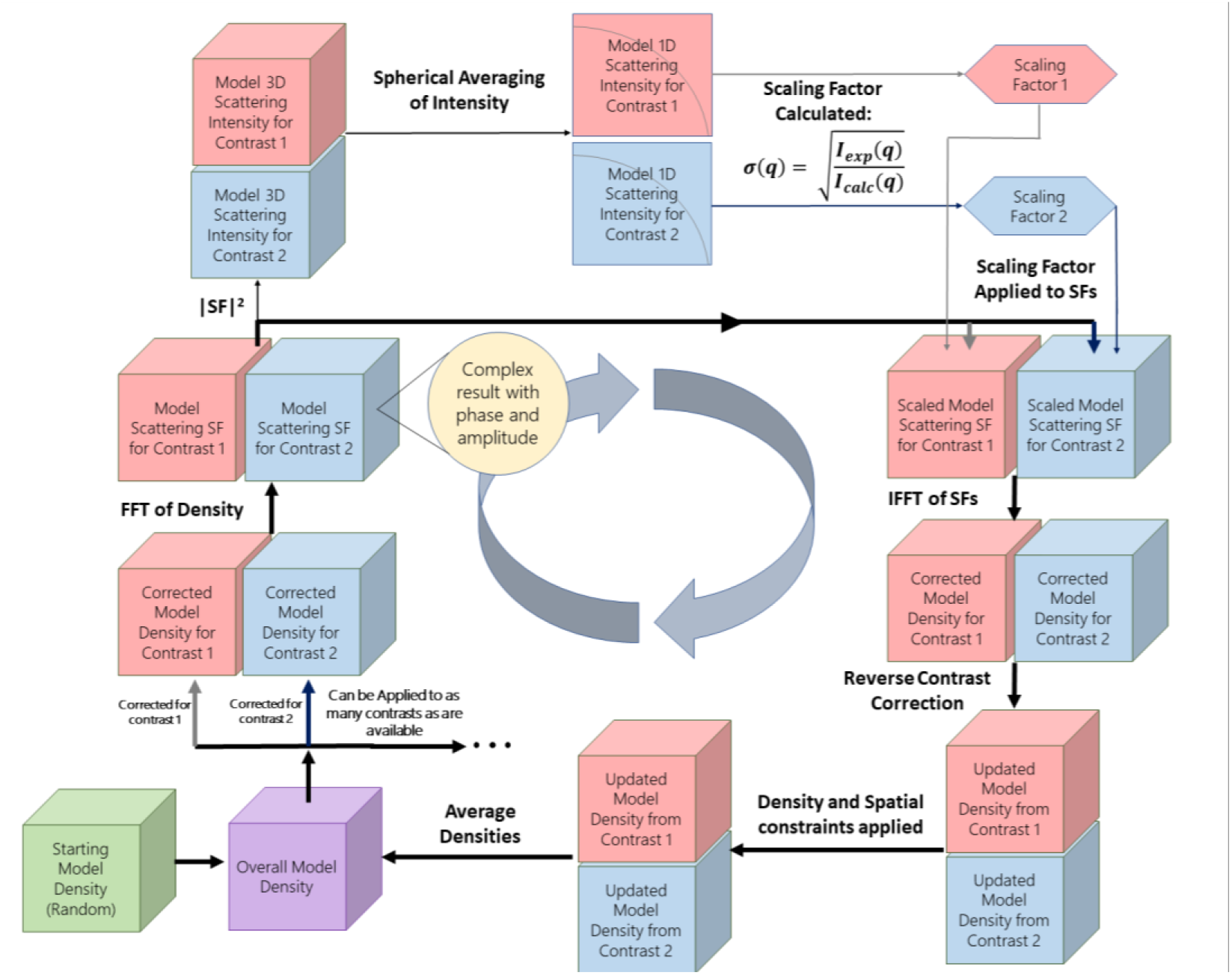
Flow chart of the DENSS-Multiple method. All structure information from different SANS contrasts is combined during iterations.

One of the most important considerations of this algorithm was combining the resultant model densities from each contrast in the iterations in an efficient and logical manner. The most obvious of the options was a simple mathematical average across all contrasts in the 3D array. Averaging the rectified densities was a simple way to combine the unique values generated from each contrast without biasing the outcome toward a specific contrast. Also, an important factor to consider is when the averaging should occur as the process iterates. We determined that averaging the densities of each contrast should occur after the real space restraints (via the support) are applied to the densities. The support discriminates between what is and what is not part of the reconstruction and is optimized using the *shrinkwrap* algorithm.^6^ This was done to maximize the heterogeneities between different contrasts. For this reason, structure-factor averaging in Fourier space was ruled out because doing so potentially cancels out some structure factors, thereby negatively affecting reconstruction.

Another important factor to consider with averaging the densities of each contrast together is the frequency of averaging. Averaging all structures with each iteration would inherently affect the next iteration for all models, thereby diminishing possible unique features or information that each contrast could provide. However, the orientations and locations of the densities from different contrasts and iterations will drift significantly in 3D space. Averaging the densities after each step essentially enforces the alignment of densities generated in different contrasts and iterations. Otherwise, it requires an alignment algorithm that is prohibitively expensive in terms of computation, or it even results in completely losing the relative alignment.

Convergence of DENSS-Multiple now depends on the χ^2^ values from each contrast. When the average variance of the previous 100 χ^2^ values for each contrast reaches a point below the break condition, the algorithm is determined to have *converged*, and the resulting model is the reconstruction for all the given NCV data. This approach ensures a stable fit to all data sets instead of one preferential data set.

We tested DENSS-Multiple on several single-phase and multiphase systems using simulated and experimental data to assess its efficacy and performance. It has matched or outperformed SAXS and SANS data in each test case, which shows that it can reconstruct envelopes and provide insight on internal neutron SLD with a structure.

### Test Examples

We tested the ability of DENSS-Multiple to resolve simple geometric models at different neutron scattering contrasts with simulated data. We created nine models, including single-phase and multiphase systems, with NCV data sets. The single-phase system represents a particle composed of only a single macromolecule class, such as a protein. The structure is usually well represented by one SAS profile; however, contrast variation provides additional data and increases the reliability of the result. The single-phase SANS contrasts were tested with various simulated and experimental data sets (Figures S1–S4). The results improved slightly from a single SANS curve, especially on structures with more complexity, such as a hollow sphere (Figure S2b).

Of particular interest were systems with multiple phases, such as particles with regions of vastly different densities resembling real complex systems (e.g., lipid nanodiscs, protein-DNA complex). The multiphase DENSS-Multiple tests are presented in Figures 2, S5, and S6 with various simulated data sets. DENSS-Multiple yielded well-defined reconstructions for almost all cases (Figures S5A, S5C, and S5D). This is of particular interest because the multiphase nature of the structure is represented by the corresponding phase contrast in the reconstructions.

**Figure 2:**
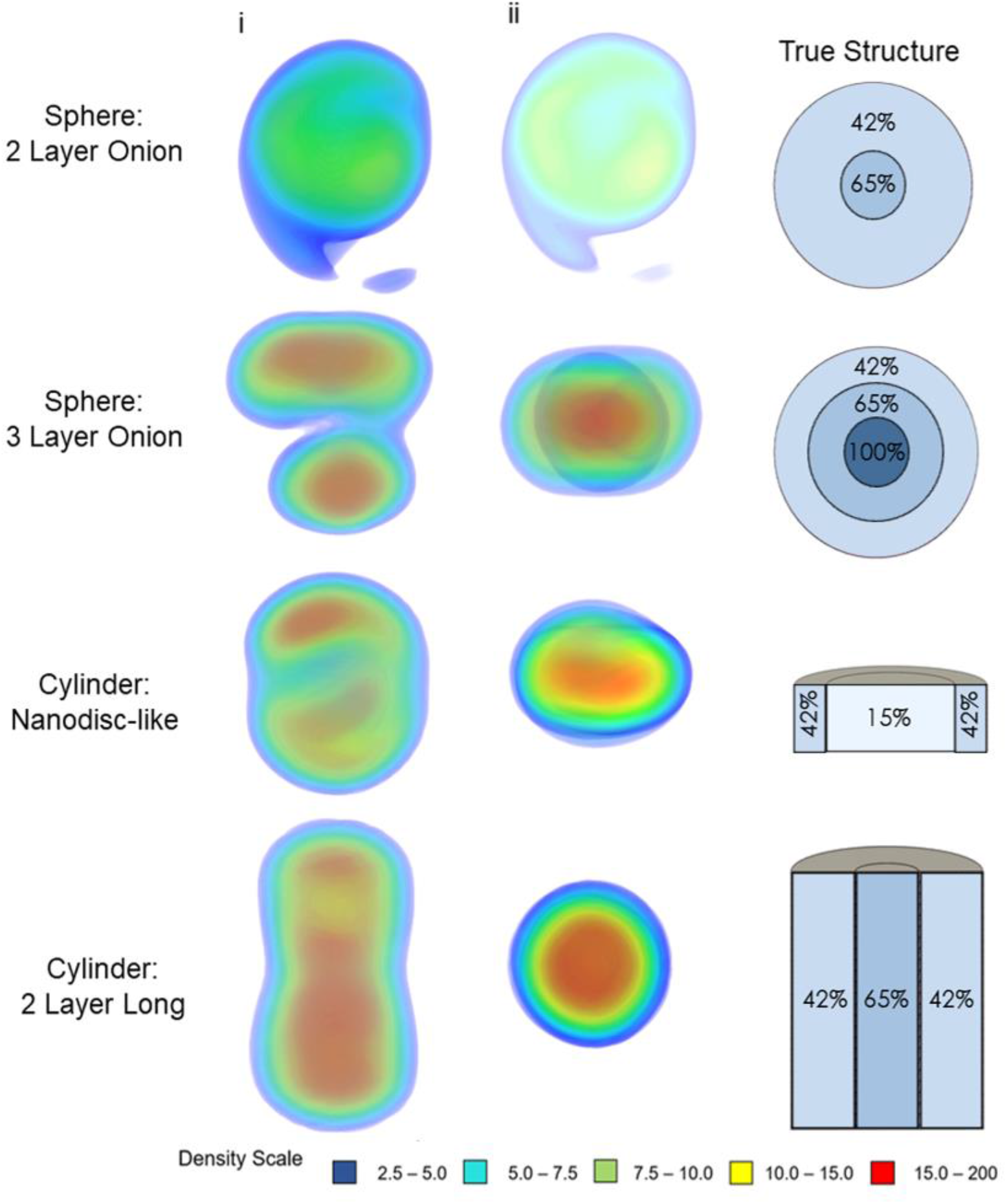
Multiphase geometric testing of DENSS-Multiple. The DENSS-Multiple results (i and ii) and the original mode structures are shown. The diameter of all simulated objects is 150 Å. All samples were analyzed and observed using PyMOL.^7^ The first shape tested was a sphere with two density layers. The sphere had an overall radius of 75 Å; the innermost core sphere had a neutron scattering length density (NSLD) equivalent to 65% D_2_O and a radius of 25 Å, and the outermost density layer had a density equivalent to 42% D_2_O and a layer thickness of 50 Å. The data was generated with background SLD values equivalent to 0%, 65%, and 100% D_2_O concentrations.

Test results with a three-layer onion model (Figures 2 and S5b) have shown that DENSS-Multiple had some difficulty using all the NCV data to generate a good result and highlighted an important point to consider when using NCV data. In a typical SANS NCV experiment, some contrasts are often designed to be cancelled out a certain phase completely (contrast matching).

A whole structure complex can be solved piece by piece and then be placed together using methods such as rigid modeling or molecular docking that can be constrained by an SAS data set with the overall structure. The complete contrast-matching of data results in a portion of the total structure being absent, thereby making it a more difficult test case for DENSS-Multiple. For example, in the three-layer onion model with the component contrast matching point (CMP) equivalent to 42%, 65%, and 100% D_2_O, using NCV data collected for 0%, 65%, and 100% D_2_O yielded results that are not representative of the starting structure. Because the NCV coincides with a significant number of the phases (two out of three), aligning the subphases among different contrasts during the iterations becomes very challenging. However, if complete contrast matching occurs in just one or none of the phases, like all other cases in Figure 2, then DENSS-Multiple is quite effective. Therefore, the use of complete contrast matching of phases should be carefully considered when using DENSS-Multiple. More tests and comments can be found in the Supplemental Materials and Figure S7.

An experimental data set with a lipid nanodisc^8^ made with membrane scaffold protein (MSP) was used for the test. The lipid nanodisc consisted of cNW9 MSP with chain-deuterated d54-DMPC and DMPC lipid in a ratio of 1.65:1, in which the averaged lipid NSLD was ~60% D_2_O. The cNW9 nanodisc’s experimental SANS NCV series consisted of 4 contrasts: 0%, 42%, 80%, and 100%. The DENSS-Multiple reconstruction of the lipid nanodisc had a structure with noticeably less noise than the single DENSS reconstructions (Figure 3). Additionally, the resolution for the DENSS-Multiple reconstruction (based on the value at which Fourier shell correlation (FSC) falls under 0.5) is better (22.3 Å) compared with the single DENSS reconstructions (~30–40 Å). This indicates that the DENSS-Multiple reconstructions combine information from different contrasts effectively and consistently to highlight different parts of the overall structure and subsequently improve the result. Additional scattering curves from various contrasts provide more constraints on reconstruction and potentially cancel out noise from different measurements.

**Figure 3:**
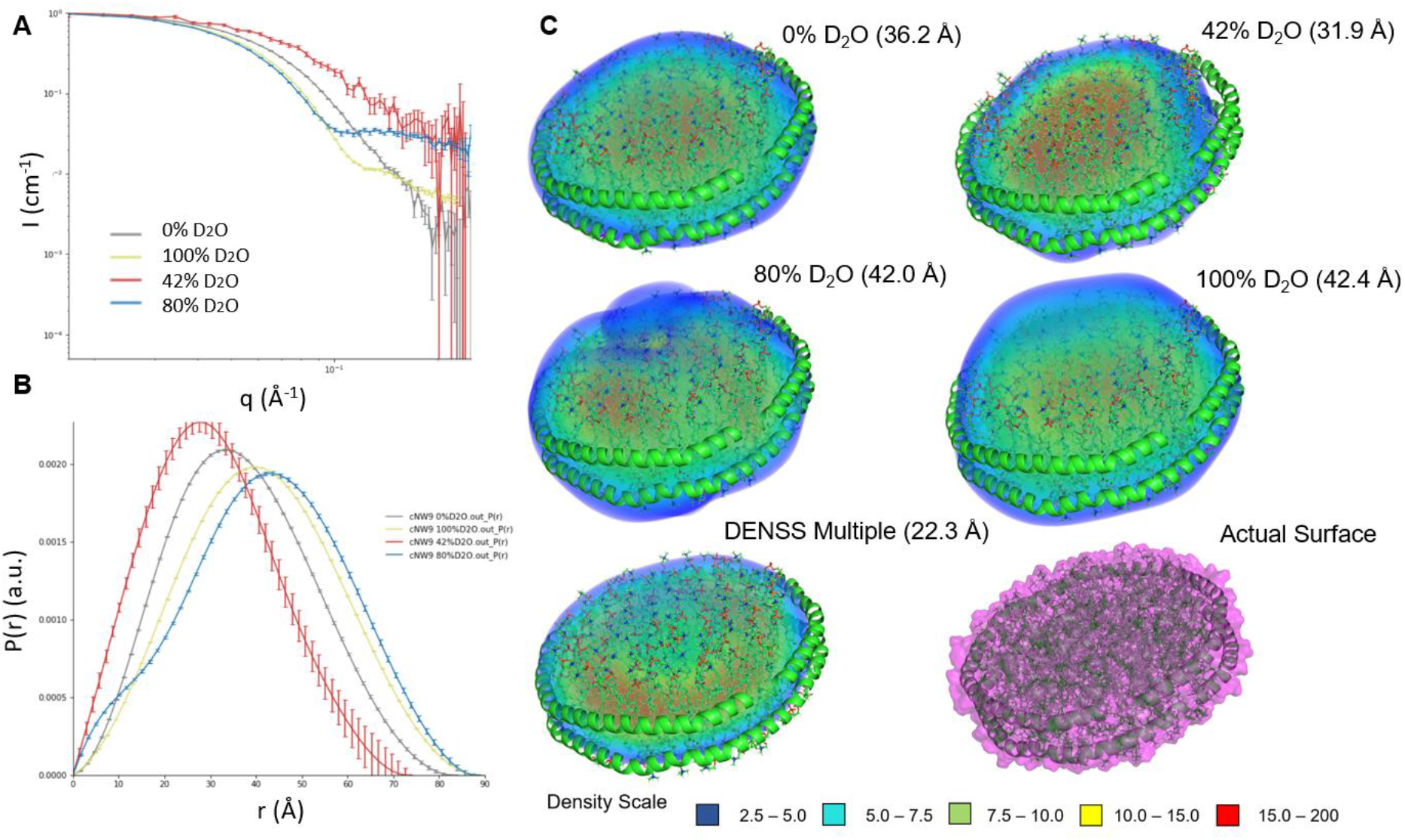
cNW9 lipid nanodisc SANS NCV series and DENSS-Multiple reconstruction. (A) SANS NCV scattering curves of CytC, shown on a log(x)-log(y) scale with error bars shown. (B) P(r) curves for scattering data in (a), generated using GNOM. (C) Normalized and averaged density maps generated by DENSS and DENSS-Multiple with calculated FSC resolution shown in parentheses. The DENSS-Multiple reconstruction was created using the 0%, 42%, 80%, and 100% SANS data combined.

We further tested DENSS-Multiple for its ability to resolve the absolute value of neutron SLD in a multiphase system.^5^ Using buffer NSLD values as input and SANS scattering profiles in absolute scale of the scattering cross section (cm^-1^) for each contrast, we found that the different phases can be discriminated easily from the final result and the density distribution histogram (Figure 4). However, the program’s output density values must be manually rescaled to achieve realistic NSLD values. We tested this approach using a three-layer onion model with phases equivalent to 100%, 65%, and 42% D_2_O from inside to outside and layer thicknesses of 25 Å each, which corresponds to a 150 Å diameter sphere. NCV data was simulated for 0%, 10%, and 20% D_2_O contrasts and reconstructed using DENSS-Multiple. The three phases are easily distinguished (Figure 4). On the density histogram, the phases are represented by the value ranges of ~0.16–0.6 for 42% D_2_O phase; 0.6–1.1 for 65% D_2_O phase; and 1.1–1.3 (max) for 100% D_2_O phase (Figure 4A). The frequency of density values near zero is quite high in all tests; this is likely caused by noise and the Gaussian smoothing applied by the DENSS algorithm in the final step of reconstruction.^5^ For each phase, a distribution of density values is apparent instead of a discrete distribution. We believe this is partly caused by the aforementioned Gaussian smoothing employed by DENSS and DENSS-Multiple as well as the resolution limitations of SAS. Additional tests and comments with experimental data sets from lipid nanodiscs are provided in the Supplemental Materials (Figure S8).

**Figure 4:**
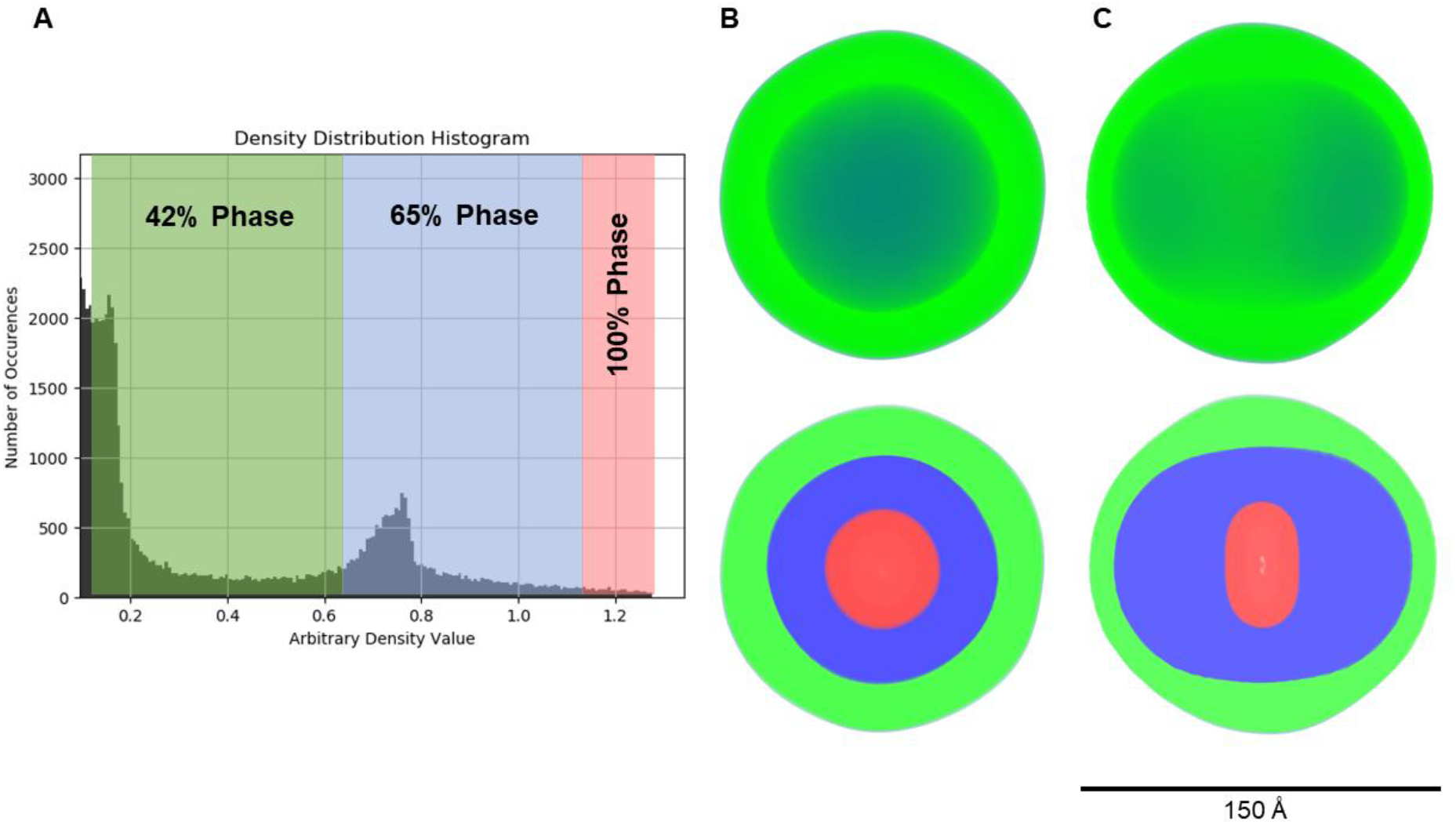
Three-layer onion sphere model reconstruction showing different phases via absolute density output from DENSS-Multiple. (A) Density distribution histogram showing the arbitrary density value output from DENSS- Multiple and the number of occurrences of that bin for each voxel of the reconstruction. Phase ranges were determined from knowledge of the three-layer onion sphere’s phase diameters. (B) Front and (C) orthogonal view of the three-layer onion sphere reconstruction (above) and slices to the center (below) with phases colored according to the ranges in (A).

## Conclusion

The significant contrast manipulation in SANS provides a great opportunity for density reconstruction with improved results compared with relatively small electron density contrast in SAXS. DENSS-Multiple is successful in accurately reconstructing the envelopes of both single- and multiphase data from simulated and experimental SANS data. This work demonstrates that DENSS-Multiple is applicable to a wide range of biomolecular systems measured with SANS NCV series methods. DENSS-Multiple and its unique ability to resolve ab initio reconstructions from multiple SANS data sets concurrently can improve reconstruction quality and resolution. We hope that others continue to iterate upon this open-source work and continue to push forward the DENSS software suite for the SAS scientific community. Currently the code is available upon request.

## Acknowledgement

We thank Qiu Zhang, Swati Pant, and Dr. Hugh O’Neill for their assistance in preparing lipid nanodisc samples for the SANS experiment.

This research was supported through the Center for Structural Molecular Biology, a user facility supported by the US Department of Energy’s (DOE’) Office of Biological and Environmental Research. This research used resources of the High Flux Isotope Reactor and the Spallation Neutron Source, both of which are DOE Office of Science User Facilities operated by the Oak Ridge National Laboratory (ORNL). Shuo Qian was partly supported by the Spallation Neutron Source Second Target Station Project at ORNL. ORNL is managed by UT-Battelle LLC for DOE’s Office of Science.

