## Supplemental Materials for "DENSS-Multiple: A Structure Reconstruction Method using Multiple Contrast Variation of Small-Angle Neutron Scattering Based on the DENSS Algorithm"


### *Experiments and Data Treatment*

All SANS data were collected at room temperature on either Bio-SANS (CG-3B) at HFIR or EQSANS (BL-6) at SNS of Oak Ridge National Laboratory. The data was reduced using facility software with background correction and normalization. All SAXS data were collected by an in-house Rigaku Bio-SAXS 2000 instrument.

The simulated SAXS or SANS data from Geometric models that can be described by analytical functions were generated in SasView 5.0 (sasview.org). All simulated protein SAXS and SANS scattering data were generated using CRYSOL and CRYSON[1], respectively. The number of points interpolated for data simulated with CRYSOL and CRYSON was 471 and 200 to mimic typical SAS data, e.g., the in-house Bio-SAXS and Bio-SANS instruments, respectively. The number of spherical harmonics used to simulate data using CRYSON and CRYSOL was dependent on the radius of gyration of the protein being simulated. Those profiles don't have error bars to represent ideal scattering intensities. All PDB models used to simulate the scattering data were obtained from the RCSB Protein Data Bank (www.RCSB.org). GNOM (ATSAS, EMBL) was used to generate P(r) distributions, the output of which was used to run DENSS and DENSS-Multiple.

### Additional Single Phase Reconstructions Test

The single-phase results (Figure S1 and S2) show that DENSS-Multiple can reliably result in more geometrically accurate reconstructions visually than single data set from DENSS and DAMMIF. The DENSS-Multiple and DAMMIF (100% $D_2O$ only) results can be seen in Figure S1, more details using DENSS are listed in Figure S2. DENSS-Multiple succeeds at reconstructing a sphere, more uniform than DENSS reconstruction in some of the contrasts showing cavity at the center (Figure S2(A)). The hollow sphere contains a 25Å radius concentric cavity providing more of a complex topology for testing (Figure S2(B)). DENSS-Multiple has an improved integrity, and more accurate cavity, e.g., the ratio of inner and outer layer is more truthful to the original model than DAMMIF results yet is less spherical similar to the original DENSS result from a single contrast. The cylinder disk is a challenge shape for any algorithm to accurately reconstruct, with no exception for DENSS-Multiple (Figure S2(C)). A cube was tested due to its sharp right angles, and DENSS-Multiple result is visually better, but still difficult to reproduce the sharpness, due to SAS resolution limit. Intuitively, DENSS-Multiple provides a combination of single DENSS results weighted by contrasts. The process benefits from increased information content provided by the contrast variation data, even in a single phase. Overall, DENSS-Multiple shows its robust ability to reconstruct shapes at least as well as, and often better than single data set reconstructions by existing algorithms. This indicates that it can be beneficial to use NCV data in ab initio reconstructions for improving single phase results.

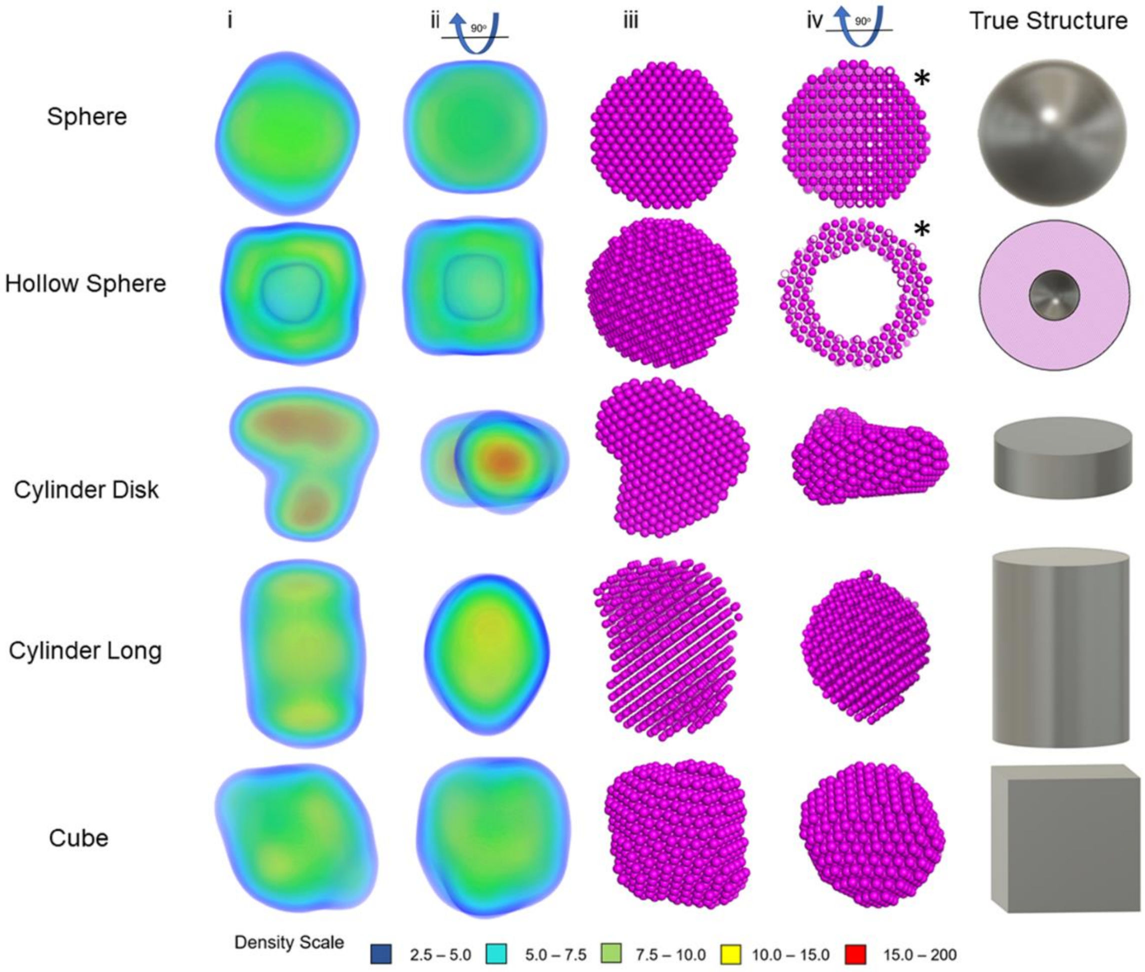


Figure S1: Single-phase geometric testing of DENSS-Multiple. Each shape tested is labeled in the left column. All shapes have a simulated NSLD of 42% $D_2O$. The DENSS-Multiple reconstructions were done using 0%, 80%, and 100% $D_2O$ data SANS contrasts, and the front view (i) and 90 degree rotation (ii) are shown, as well as the DAMMIF 100% $D_2O$ contrasts in a normal view (iii) and 90 degree rotation (iv), except for the sphere and hollow sphere (marked with asterisk), in which a central slice is shown for (iv). The structure models are shown on the right. All samples were analyzed and visualized using PyMOL [2]. Density scale is in arbitrary unit. All densities are normalized to the same scale for visualization and comparison. All models were set up to have a diameter or side length of 150Å for consistency.

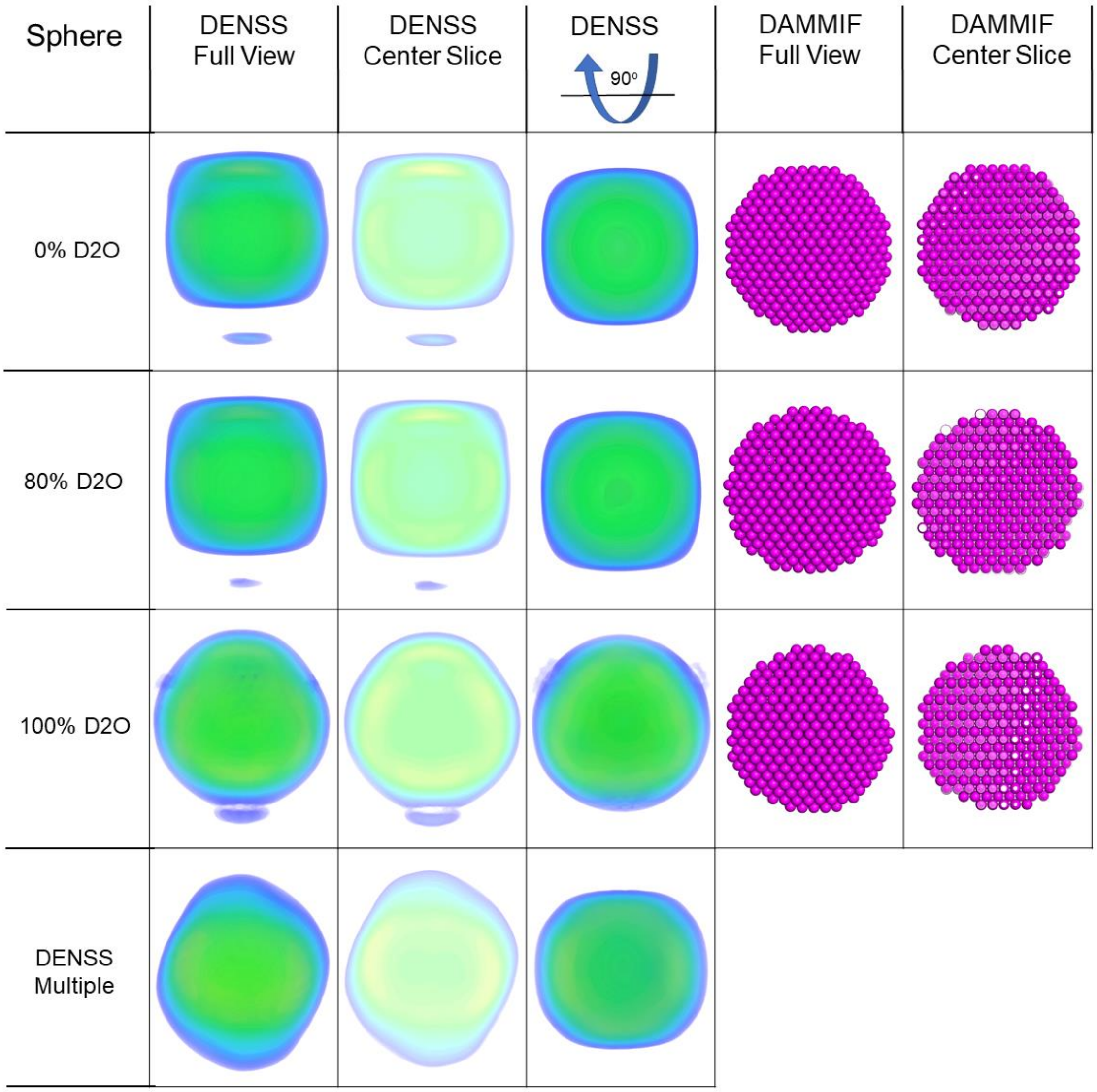


A. Shape: Sphere. The results from a single SANS contrast by DENSS/DAMMIF were compared with the result from NCV by DENSS-Multiple

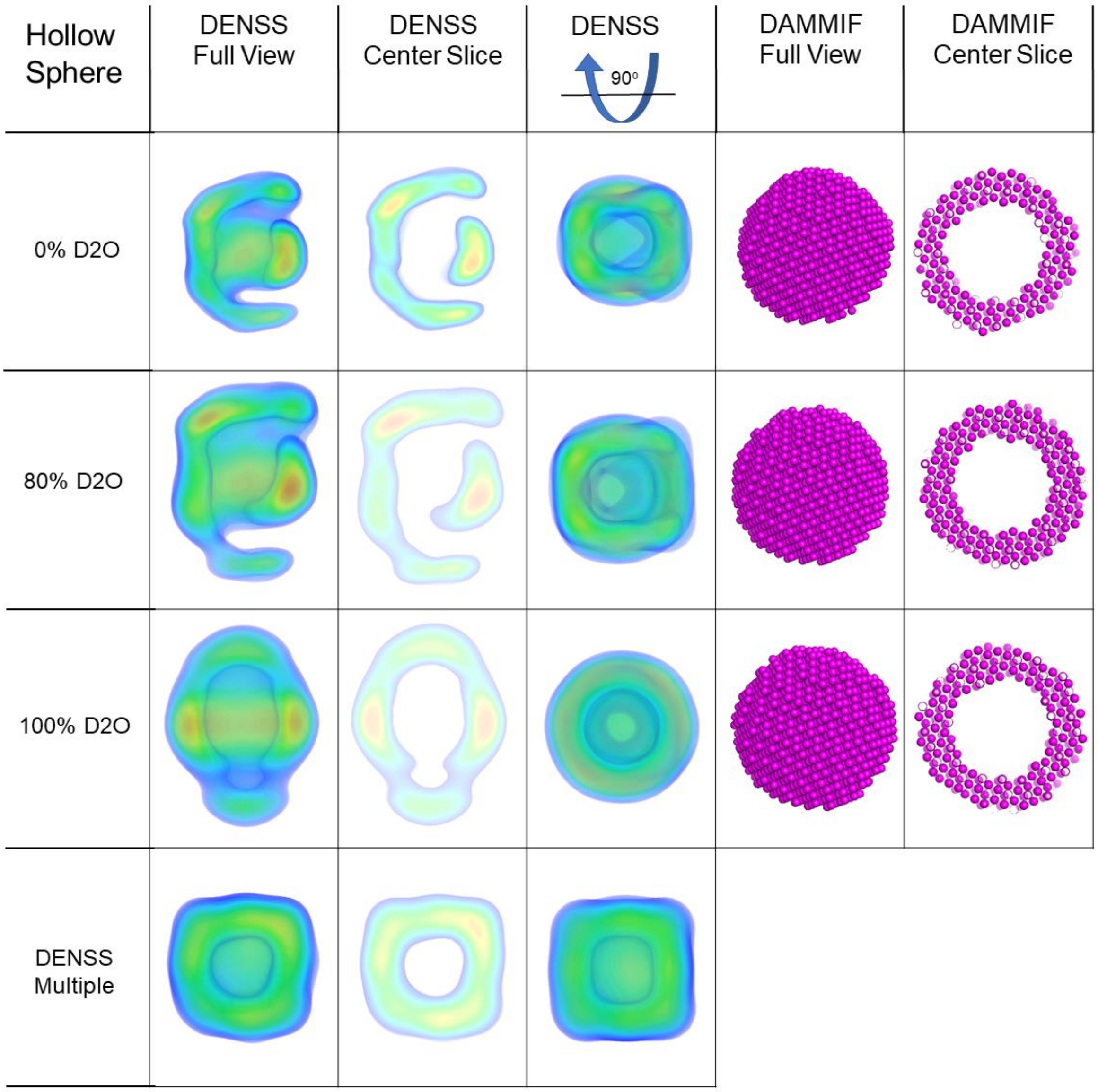


B. Shape: Hollow sphere. The results from a single SANS contrast by DENSS/DAMMIF were compared with the result from NCV by DENSS-Multiple

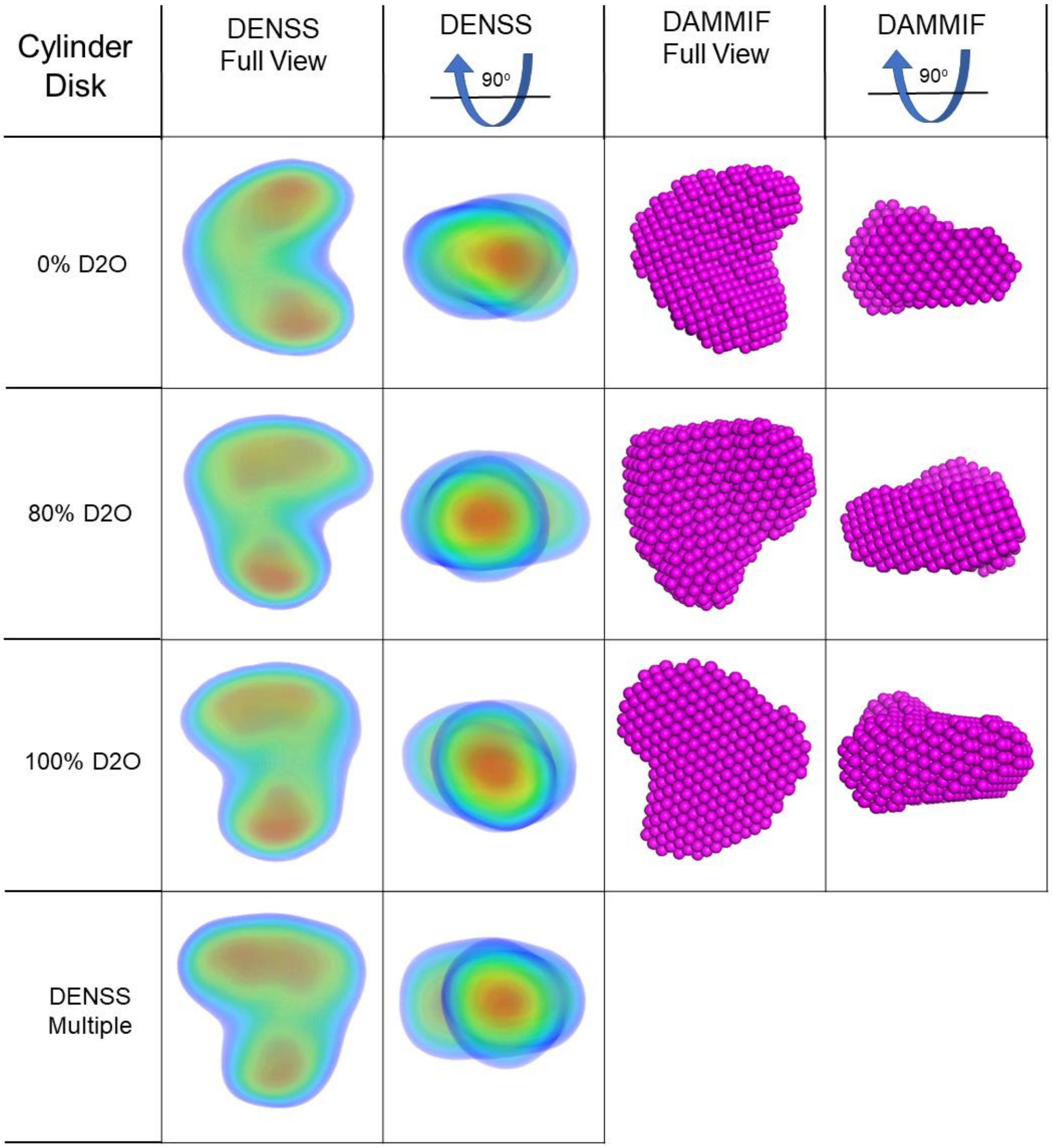


C: Shape: cylinder disk. The results from a single SANS contrast by DENSS/ DAMMIF were compared with the result from NCV by DENSS-Multiple

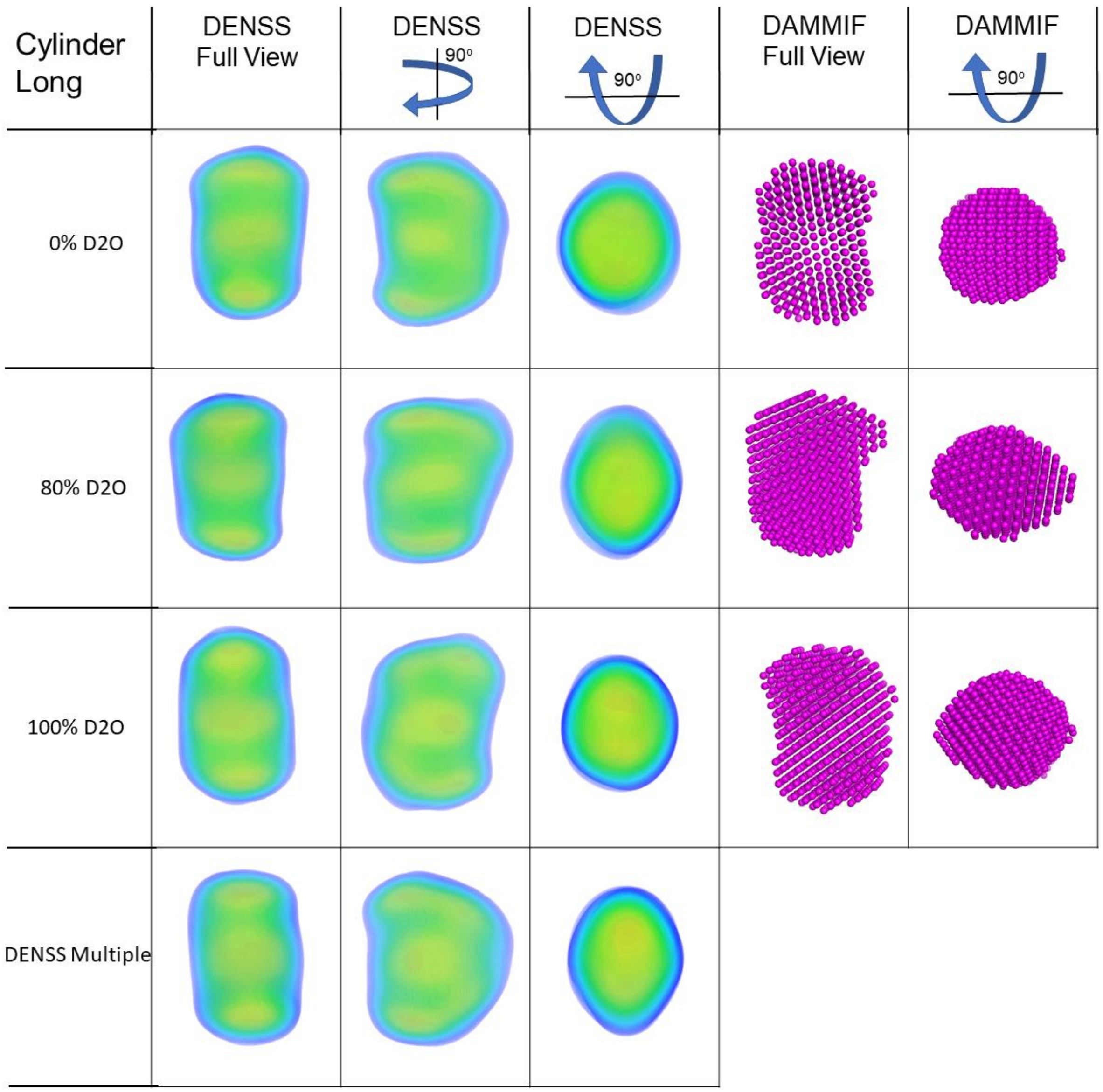


D: Shape: Long Cylinder . The results from a single SANS contrast by DENSS/ DAMMIF were compared with the result from NCV by DENSS-Multiple

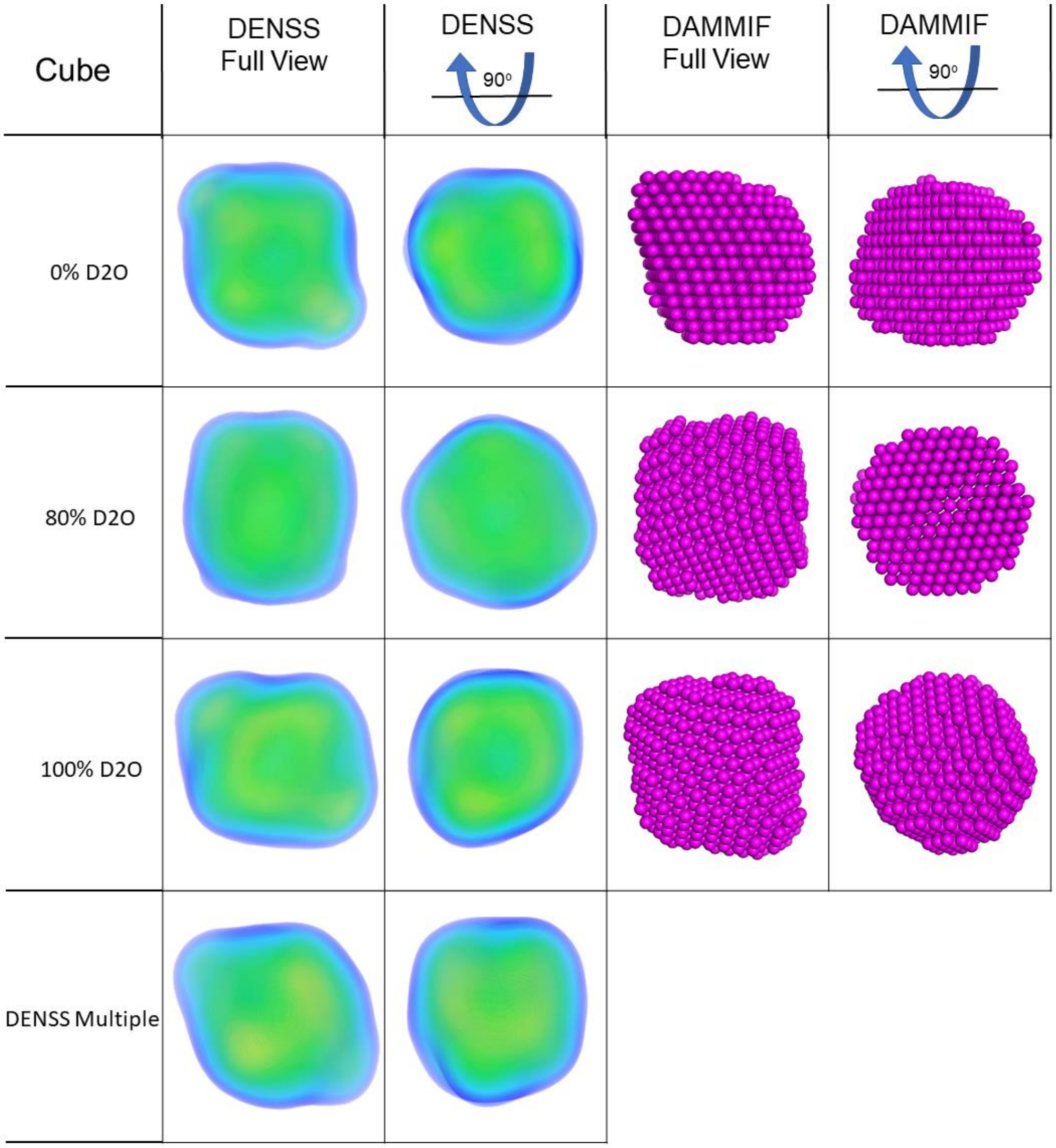


E. Shape: Cube. The results from a single SANS contrast by DENSS/DAMMIF were compared with the result from NCV by DENSS-Multiple

Figure S2: Single Phase Testing with DENSS-Multiple (A-E). DENSS and DAMMIF models shown as labeled. The shapes are labelled on the panels.

### Single-Phase Test with Experimental Data

Human serum albumin (HSA) in 0%, 20%, 60%, 80% and 100% $D_2O$ were measured for this test. The results are presented in Figure. S3 and S4. The SAXS and DENSS-Multiple reconstructions accurately represent the expected protein envelope. Note the SANS scattering curves (Figure S3 (A)) with low contrast (i.e. HSA 60% $D_2O$) have larger error bars, which cause larger error bars in P(r) (Figure S3(B)). Therefore, there is high variability in the Dmax values for the P(r) curves, but they all peak around 38 Å indicating the overall size distribution of the particle is similar across contrasts. The individual SANS reconstructions from DENSS from the 20% and 100% $D_2O$ do not closely resemble the SAXS reconstruction nor the expected morphology of the actual protein surface. However, the 0%, 60%, and 80% $D_2O$ reconstructions overlays well with the protein PDB (PDB-ID: 3JRY)[3]. Visually, the DENSS-Multiple result overlays well with the HSA PDB, and it seems comparable to the accuracy of the SAXS reconstruction in both resolution and density. More details can be seen in Figure S4. The DENSS-Multiple reconstruction visually outperforms each of the individual SANS $D_2O$ contrast reconstructions, benefitting from additional information from each contrast measurement. The single-phase test with HSA showed the benefits of DENSS-Multiple in reconstructing an accurate structure out of multiple $D_2O$ contrasts despite noisy individual contrast data.

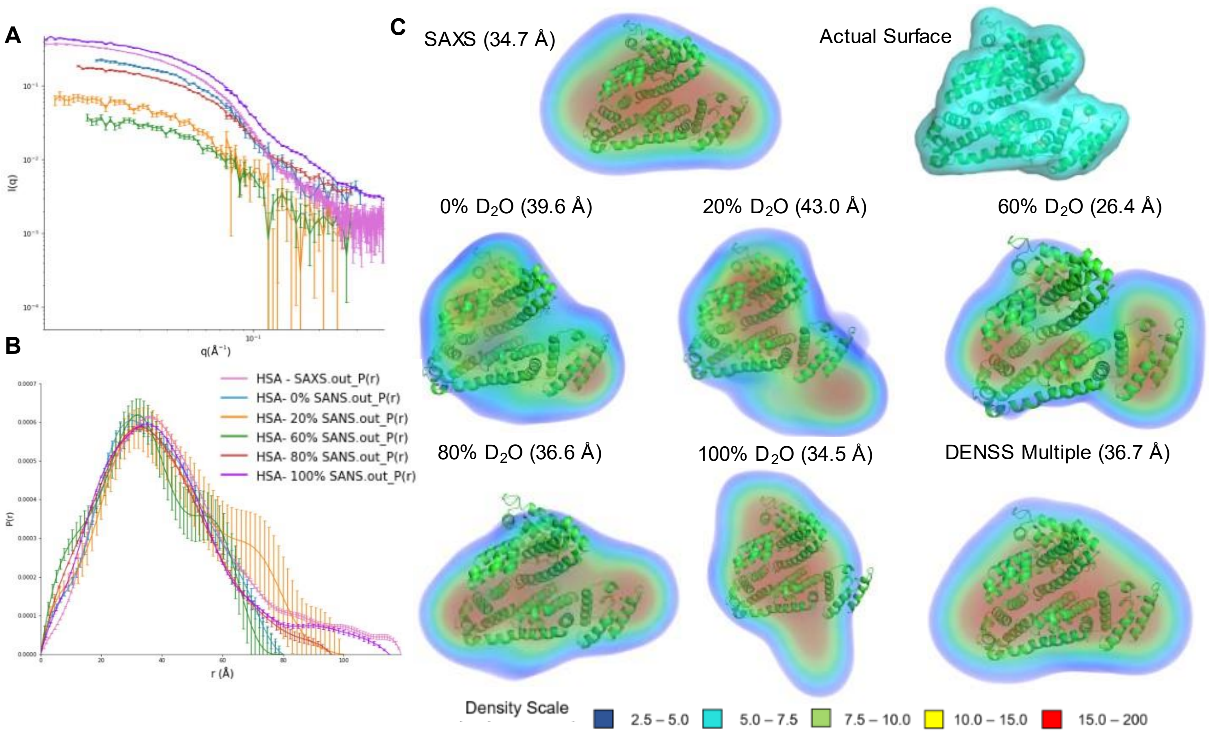


Figure S3: HSA SAXS and SANS single DENSS results compared to DENSS-Multiple. A.) SAXS and SANS scattering profiles. SAXS data was scaled down to accentuate detail of the SANS curves. B.) P(r) curves for scattering data in A, generated using GNOM. C.) Normalized density maps generated by DENSS and DENSS-Multiple. All densities are normalized to the same scale for visualization and comparison. The calculated FSC resolution is shown in parentheses. The DENSS-Multiple result was created using the 0, 20, 60, 80, and 100% $D_2O$ data. Density scale is in arbitrary unit.

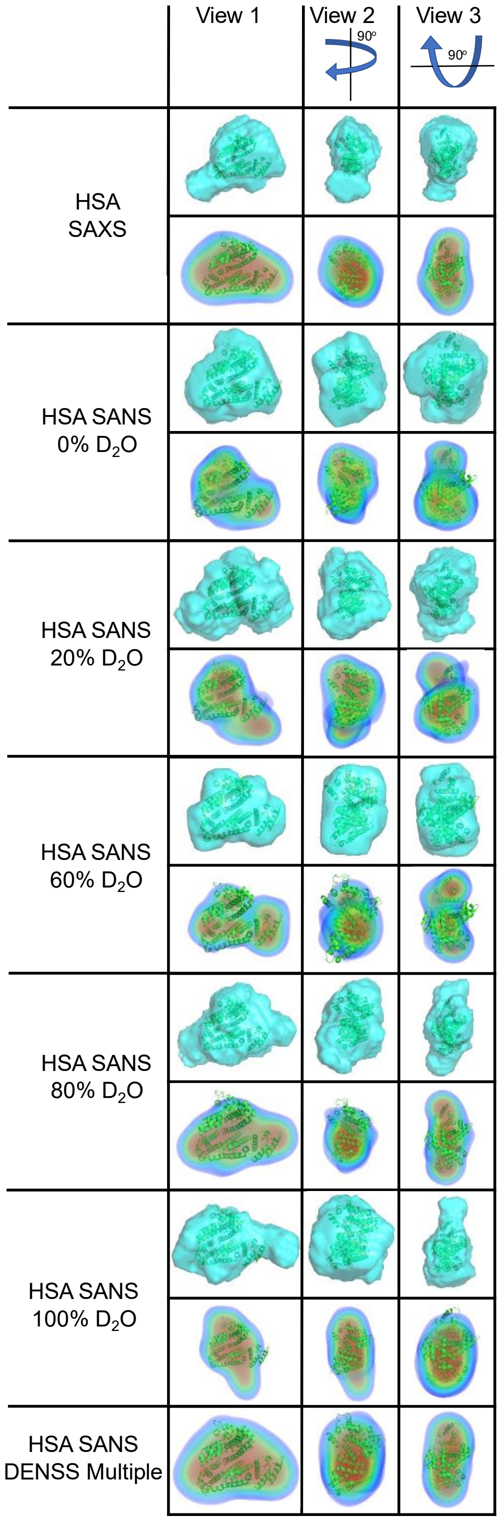


Figure S4: Human Serine Albumin (HSA) Single Phase Testing with DAMMIF (Cyan), DENSS (below DAMMIF result), and DENSS-Multiple (the final row).

**Multi-Phase Test**

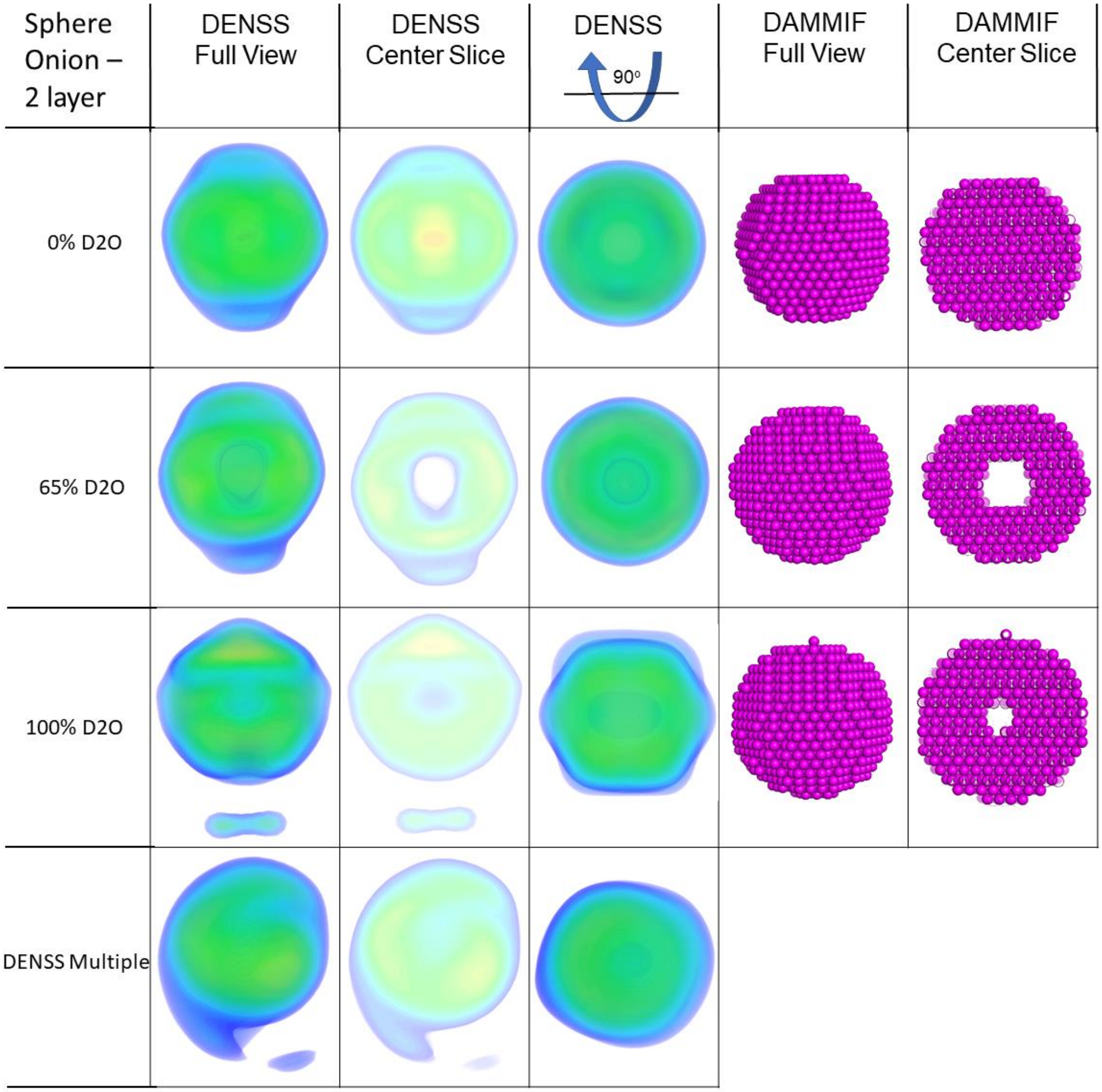


A. Shape: 2 Layer Onion Sphere. The results from a single SANS contrast by DENSS/DAMMIF were compared with the result from NCV by DENSS-Multiple

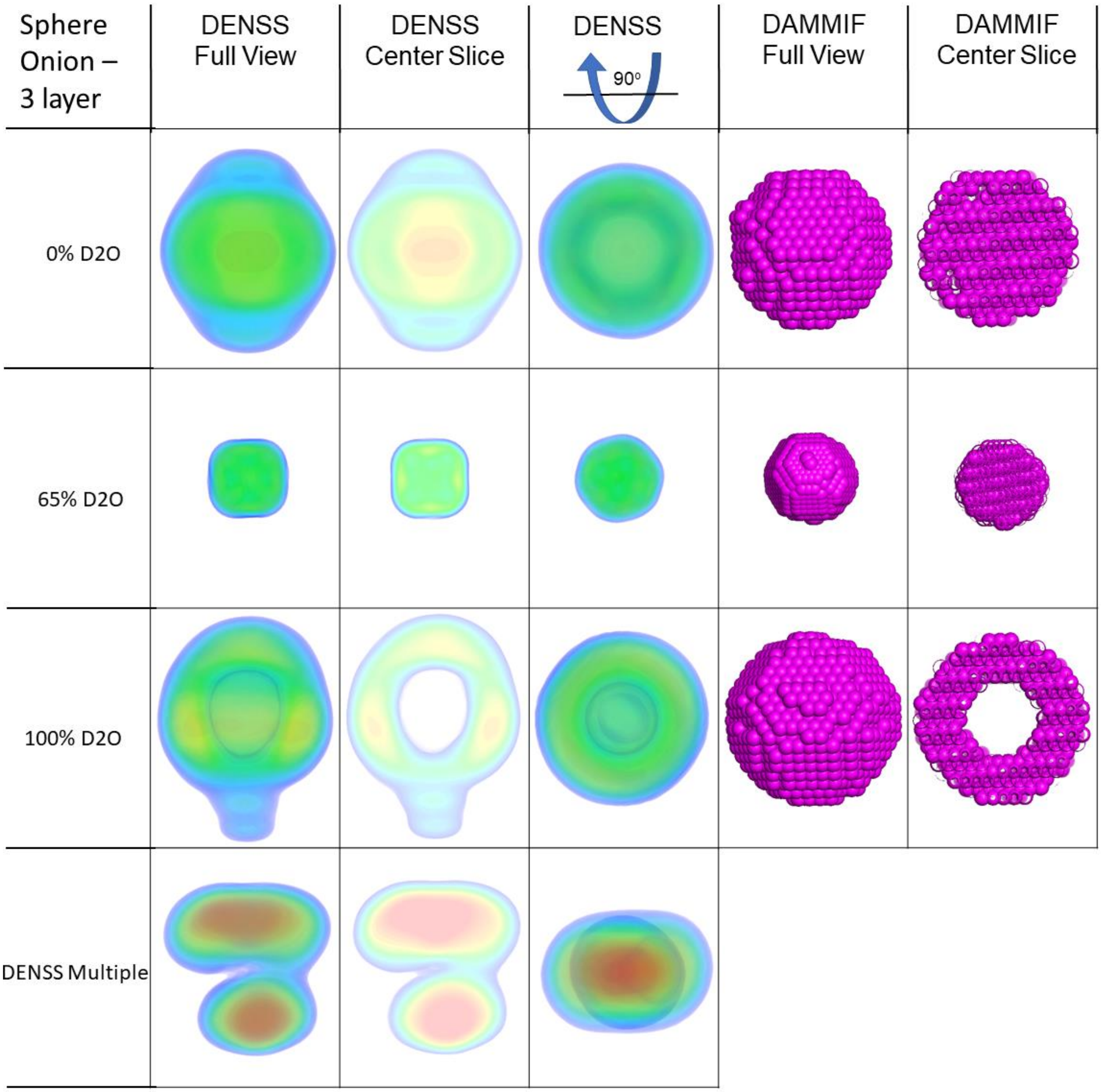


B. Shape: 3 Layer Onion Sphere. The results from a single SANS contrast by DENSS/DAMMIF were compared with the result from NCV by DENSS-Multiple

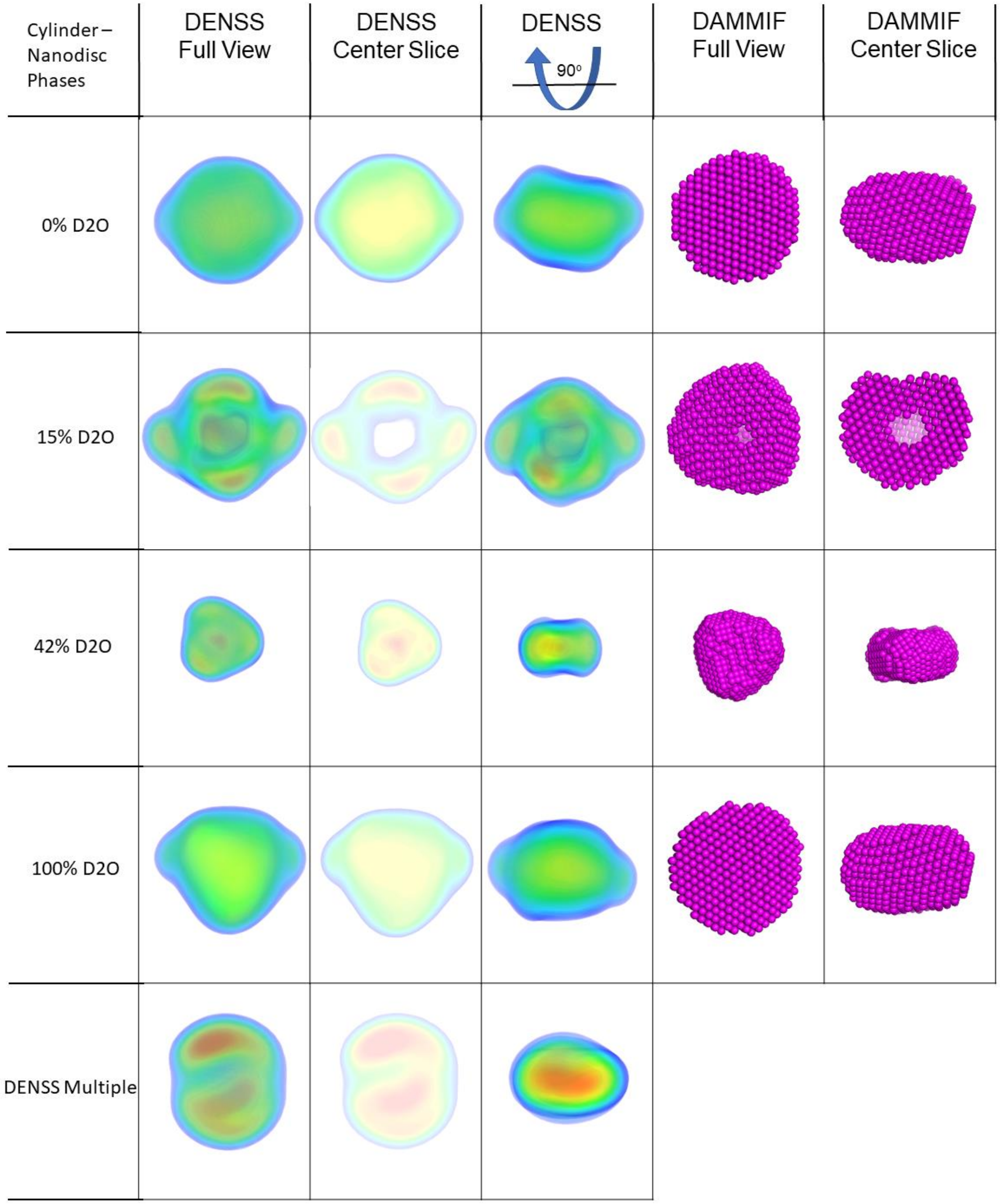


C: Shape: nanodisc-like cylinder. The results from a single SANS contrast by DENSS/DAMMIF were compared with the result from NCV by DENSS-Multiple

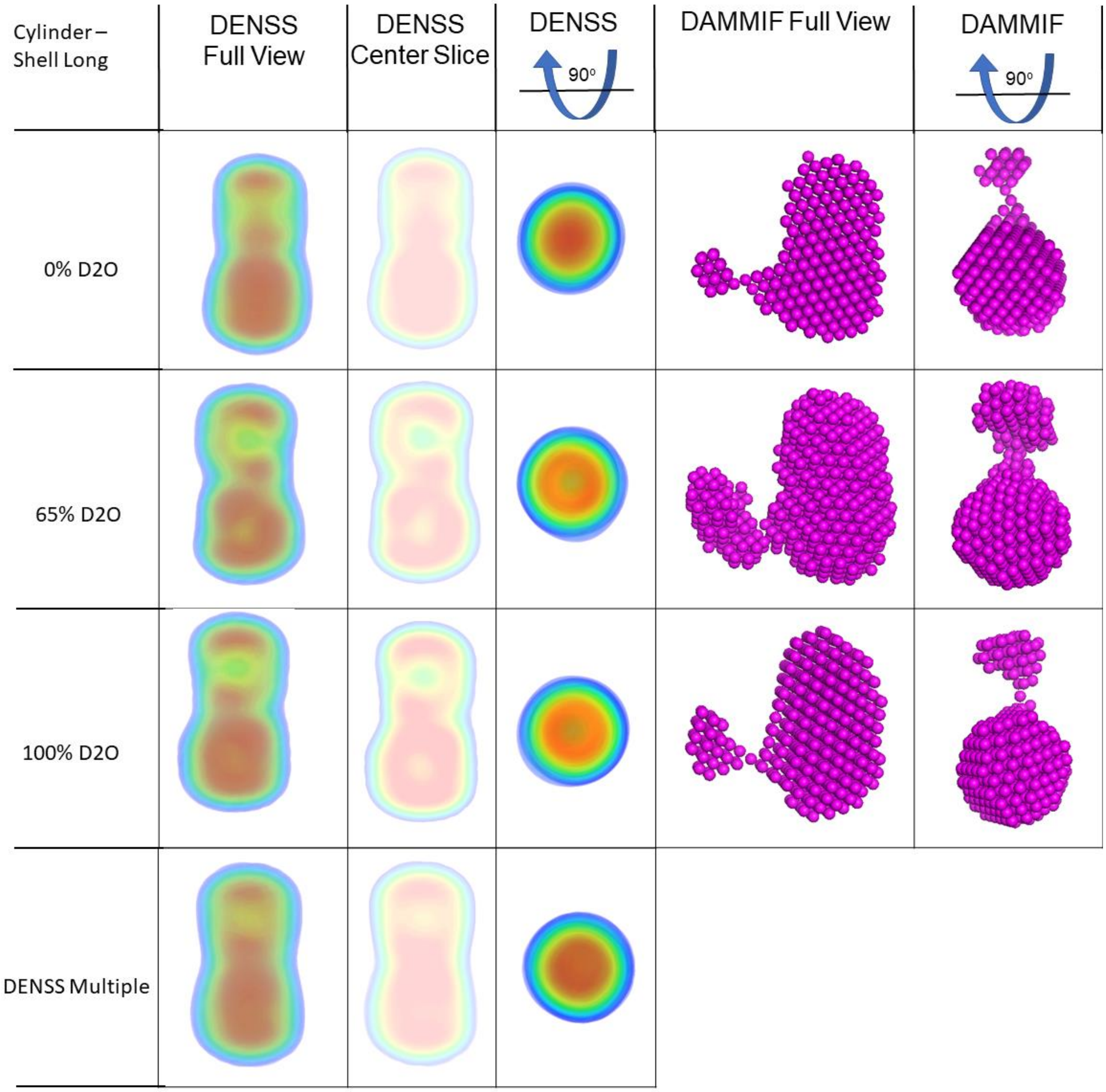


D. Shape: Cylinder with 2 layers. The results from a single SANS contrast by DENSS/DAMMIF were compared with the result from NCV by DENSS-Multiple

**Figure S5:** Multi-phase test with DENSS-Multiple (A-D). This provides more details to supplement Figure 2 in the main text.

**Additional Multi-Phase Test**

An artificial HSA dimer with separate protiated and deuterated monomers was created as a simple two-phase particle for this test. The hydrogenated monomer's contrast match point is at 42% $D_2O$ and the deuterated one is at 100% $D_2O$. Three contrasts were simulated: 0%, 42%, and 100% $D_2O$, resulting in an overall dimer contrast and one contrast for each of the dimers. The difference in the particle sizes can be seen directly from the scattering data (Figure S6 (A)), where the 0% contrast has a higher Rg (37 Å) than the 42% and 100% contrasts (28 Å and 23 Å, respectively). Similarly, the P(r) curves show that the 42% and 100% contrasts have much smaller $D_{max}$ values than the 0% (Figure S6(B)). The individual DENSS reconstructions of the SANS contrasts show a disparity in the overall size of the reconstructed particles, with both the 42% and 100% contrasts reconstructing an envelope the size of the HSA monomer, while the 0% reconstruction reconstructed an envelope which fit best to the HSA dimer. Interestingly, the DENSS-Multiple reconstruction aligns well with the dimer yet appears noticeably smaller than the 0% reconstruction but with better resolution. It is important that DENSS-Multiple always has at least one data set shows the entire extent of the molecule to provide the overall envelope for the particle.

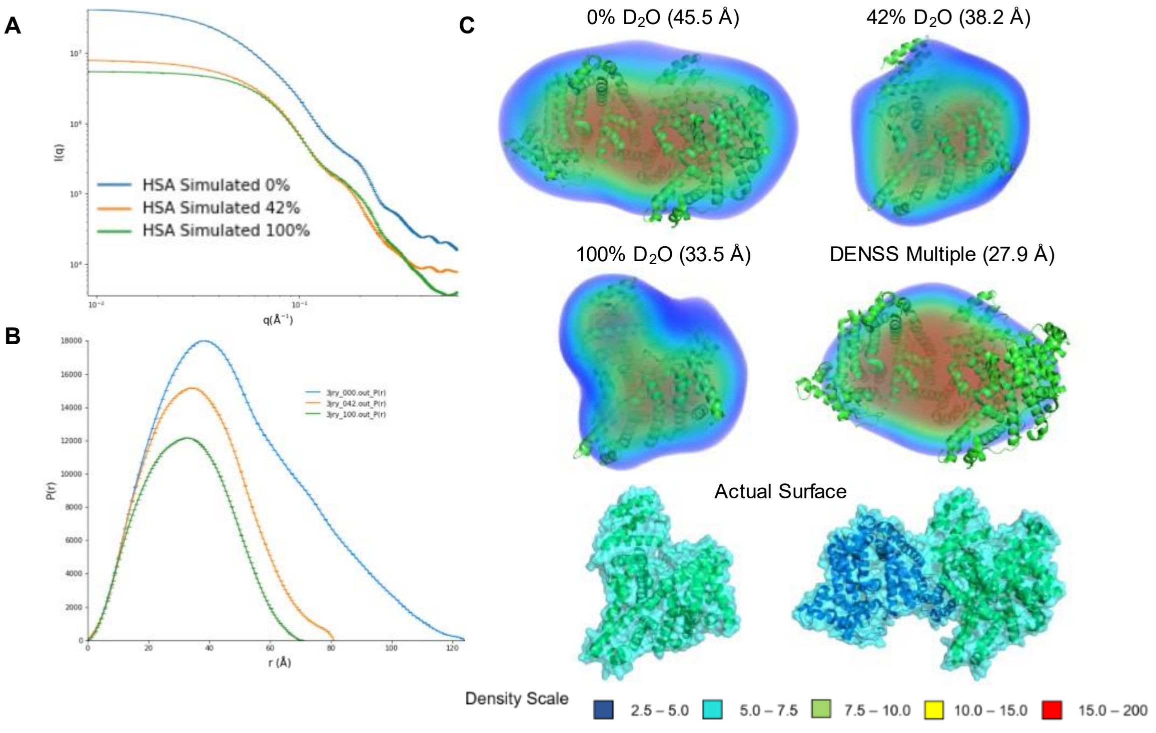


Figure S6: HSA NCV Series Simulated SANS Results Compared to DENSS-Multiple. A.) SANS Scattering curves of HSA simulated with CRYSON. B.) P(r) curves for scattering data in A, generated using GNOM. C.) Normalized density maps generated by DENSS and DENSS-Multiple. The calculated FSC resolution is shown in parentheses. The 42% and 100% $D_2O$ reconstructions are both aligned with the HSA monomer. The 0% and DENSS-Multiple reconstructions were both aligned with the HSA dimer. The DENSS-Multiple reconstruction was created using the 0%, 42%, and 100% $D_2O$ data. All densities are normalized to the same scale for visualization and comparison. Density scale is in arbitrary unit.

### Test on Contrast Inclusion

Here we demonstrate that the use of different contrast needs to be considered carefully especially for those contrasts that match out a component completely. An example of this is shown in Figure S7 with a previously test model:  2 layer onion sphere. This artificial shape has an overall diameter of 150 Å and two phases: the outer layer phase of thickness 50 Å has a SLD with CMP at 42% $D_2O$; the inner core has a SLD with CMP at 65% $D_2O$, and has a diameter of 50 Å. With such a system, the outer shell will be completely invisible in 42% $D_2O$, leaving only the inner sphere. The reconstruction issues are best seen in Figure S7, where DENSS-Multiple reconstruction results in a shape that does not resemble the sphere (i and ii) the data in 0%, 20%, and 42% $D_2O$ (Figure S7(A)). However, if the 42% $D_2O$ data is not used, thereby running DENSS-Multiple with only the 0% and 20% contrasts, the shape with 2 phases can be reconstructed successfully. Tests with 0%, 42% and 65% $D_2O$ compared to just using 0%, 65% $D_2O$ (Figure S7(B)); or with 42%, 65%, and 100% $D_2O$ compared to just using 65%, 100% $D_2O$ yield similar results (Figure S7 (C)).

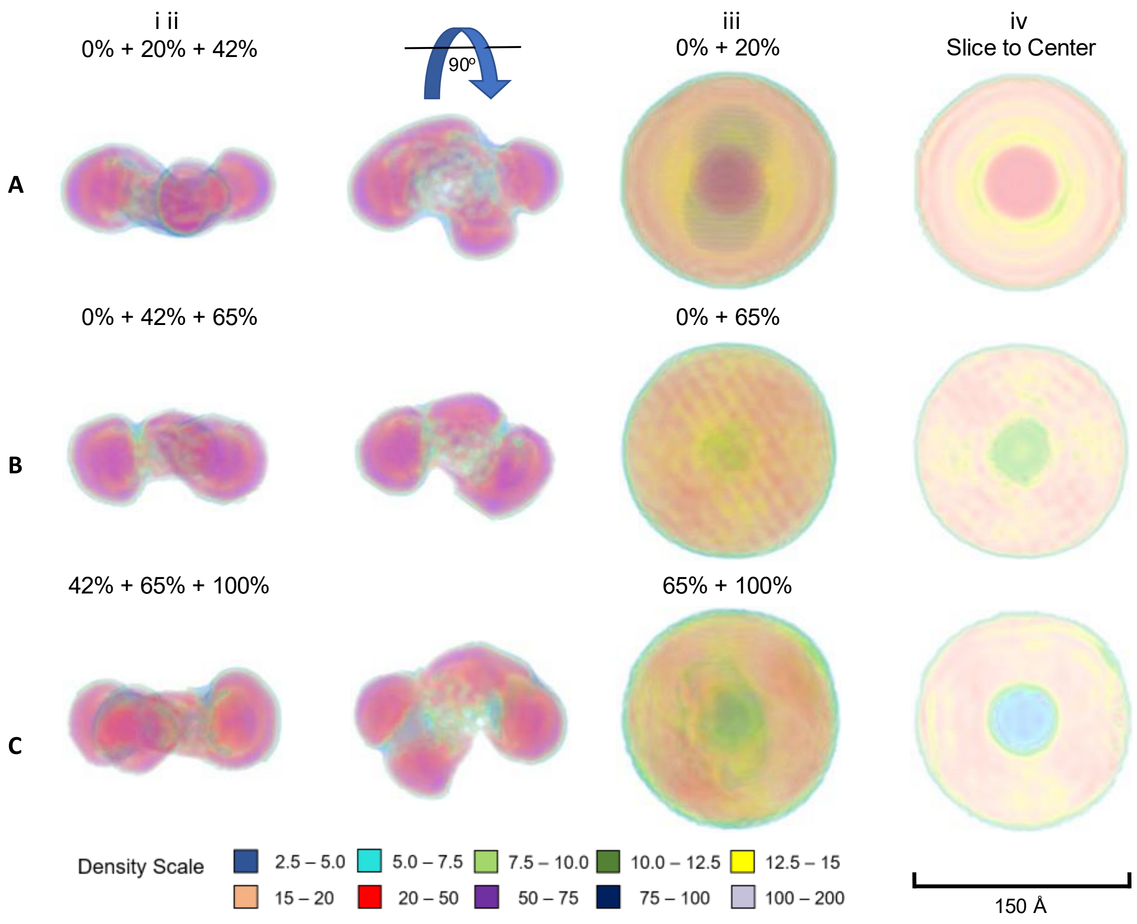


Figure S7: All reconstructions of the '2 layer onion sphere' were created using DENSS-Multiple and are shown with the density maps normalized. A) Positive contrast test with 0%, 20%, and 42% $D_2O$ contrasts (i and ii[90º rotation]) or just the 0% and 20% $D_2O$ contrasts (iii and iv[slice to center]). B) Mixed contrast test with 0%, 42% and 65% $D_2O$ contrasts (i and ii [90º rotation]) or the 0% and 65% $D_2O$ contrasts (iii and iv [slice to center]). C) Negative contrast test with 42%, 65%, and 100% $D_2O$ contrasts (i and ii[90º rotation]) or the 65% and 100% $D_2O$ contrasts (iii and iv[slice to center]). Density visualization using PyMOL. All densities are normalized to the same scale for visualization and comparison. Density scale is in arbitrary unit.

**Density Value from Different Phases**

To test whether DENSS-Multiple could accurately resolve the absolute value of neutron SLDs from NCV series data, two experimental datasets were collected from two nanodisc models. One lipid nanodisc has a protiated, circularized MSP protein with mixture of d54-DMPC/DMGP as described in the main text. The other nanodisc contained only deuterated d54-DMPC phospholipids with its tail CMP above 100% $D_2O$ and its headgroup CMP at 35% $D_2O$, and the MSP protein was partially deuterated to have the same CMP ~80% D2O. The natural variation of SLD between different components inside the nanodiscs make them good test for the effectiveness of resolving the SLD from multiple phases by DENSS-Multiple. The deuterated nanodisc was measured in ~0%, 20%, 40%, 60% and 100% $D_2O$.

The DENSS-Multiple results for both the deuterated and protiated cNW9 nanodiscs (Figure S8) were successful in shape, with DENSS-Multiple  reconstructing a discoidal density map in both cases that closely resembles the dimensions of a generated cNW9 nanodisc model and fits well to the experimental scattering data. However, as discussed in the main text, the absolute SLD in the results is not achieved. Close examination of both structures and their density distribution histogram reveals some interesting aspects. First, since the maximum SLD of the deuterated nanodisc (the deuterated tail SLD 6.83E-6 $Å^2$) is much higher the lipid part of the protiated nanodisc (CMP ~60%, SLD 3.59E-6 $Å^2$). Secondly, the highest density value is indeed much higher in the results. The much higher density in the central region of the nanodisc is well presented. For the protiated nanodisc, there is a low-density spot within the overall higher density in the lipid region. This could be an artifact. Overall, the average density for the volume is 0.03753 and 0.02894 for the deuterated and protiated nanodisc, respectively, indicating DENSS-Multiple can resolve and scale different phases qualitatively.

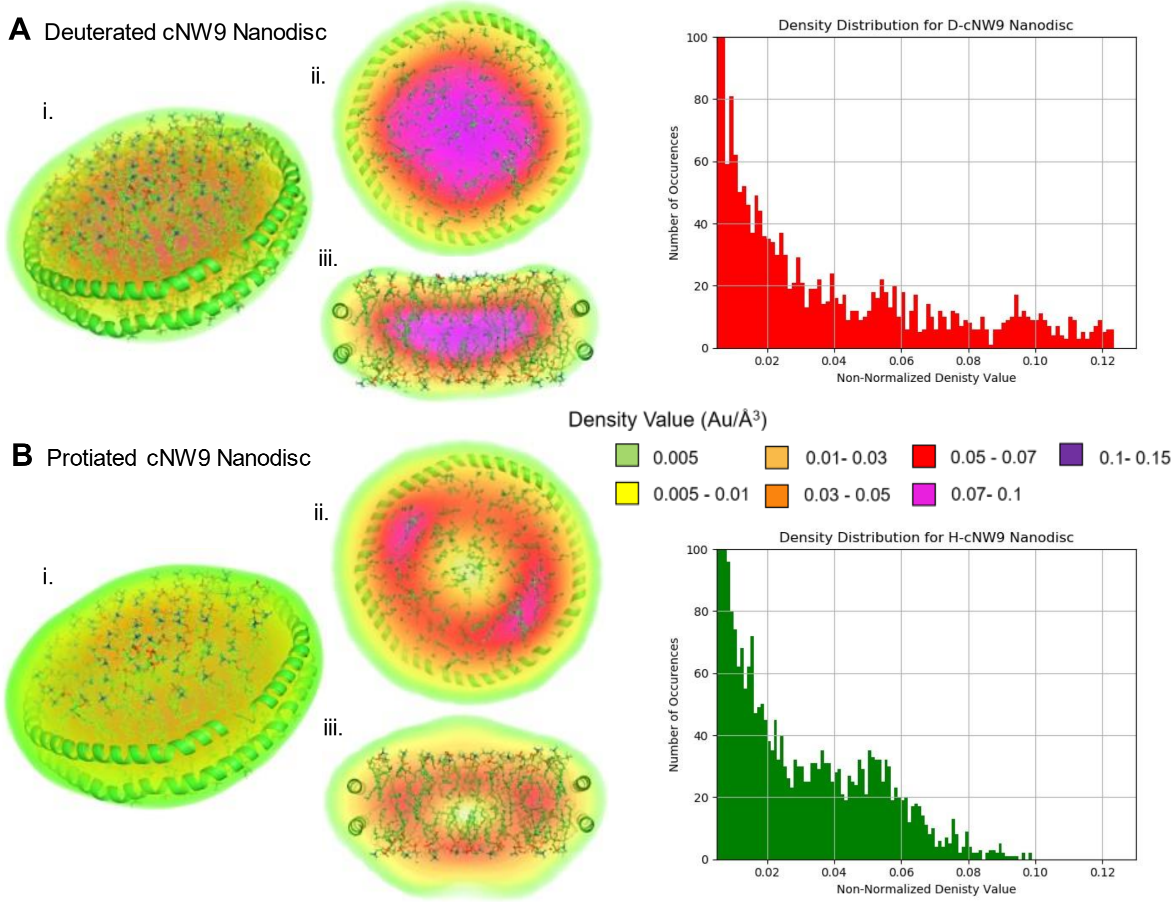


Figure S8: (A) Deuterated and (B) Protiated cNW9 nanodisc with corresponding density distribution histogram of density shown in reconstruction. Density Histograms created using Matplotlib [4,5]. i.) Diagonal view showing the full extent of the nanodisc and reconstructed density; ii.) Horizontal slice along the mid-plane of the nanodisc to show internal density; iii.) Vertical slice along the mid-plane of the nanodisc to show internal density. Color scale is in arbitrary units but the density here are not normalized between two structures and native density values from reconstructions are shown.
